# *Pyrenophora teres* f. *teres* uses specialized infection structures to facilitate effector-driven promotion of *in planta* colonization on barley

**DOI:** 10.64898/2026.11.18.767561

**Authors:** Ashley C. Nelson, Ryan M. Skiba, Michele C. Malvestiti, Gayan Kariyawasam, Pawel Borowicz, Richard D. Horsley, Zhaohui Liu, Robert S. Brueggeman, Timothy L. Friesen

## Abstract

*Pyrenophora teres* f. *teres* is a necrotrophic fungal pathogen that causes net form net blotch (NFNB) of barley. Infection of susceptible barley leaves initially causes dot-like lesions that progress to transverse and longitudinal striations, that form the net-like pattern of necrosis for which the disease was named. While the complex genetic architecture of the *P. teres* f. *teres*-barley interaction involves both dominant resistance and susceptibility and quantitative resistance, the *in planta* infection structures, compatible infection timeline, and the influence of effectors remain poorly defined. This study used fluorescent staining and confocal laser scanning microscopy (CLSM) to define infection structures and characterize the compatible infection timeline of *P. teres* f. *teres*. Using CLSM in conjunction with image-segmentation machine learning, we generated a clear comparative infection timeline showing how *P. teres* f. *teres* isolates harboring zero, one, or two virulence effector alleles infect the barley cultivar Rika. We believe that the effector genes *VR1* and *VR2* contributed to virulence in a synergistic manner. The effector genes’ combined presence accelerated the timing and spread of programmed cell death, resulting in significantly greater hyphal volume and visibly more severe macroscopic lesion development. This suggests that *P. teres* f. *teres* uses specialized infection structures alongside effectors to maximize fungal colonization and hyphal positioning to optimize nutrient uptake, perpetuating its polycyclic life cycle.

## Introduction

*Pyrenophora teres* f. *teres* is a necrotrophic fungal pathogen that causes net form net blotch (NFNB), a destructive foliar disease of barley worldwide (Shipton et al. 1973; Liu et al. 2011). Infection of susceptible barley leaves results in dot-like lesions that progress to transverse and longitudinal striations, which then form the net-like pattern of necrosis for which the disease was named (Mathre 1997). Previous microscopic and ultrastructural analyses have broadly outlined the infection process (Keeling and Banttari, 1975; Van Caeseele and Grumbles, 1979; Keon and Hargreaves, 1983), laying the groundwork for deeper investigation.

After inoculation, a conidium germinates on the leaf surface, then forms an appressorium that facilitates direct penetration of epidermal cells. Following penetration, a primary intracellular vesicle (Keeling and Banttari, 1975; Van Caeseele and Grumbles, 1979; Keon and Hargreaves, 1983), and occasionally a secondary vesicle (Keon and Hargreaves, 1983), develop, from which two to three bulbous intracellular hyphae emerge (Keeling and Banttari, 1975; Keon and Hargreaves, 1983). These hyphae eventually exit the epidermal layer to penetrate and colonize the mesophyll layer intercellularly, ultimately leading to cell death and degradation of the barley leaf (Keeling and Banttari, 1975; Van Caeseele and Grumbles, 1979; Keon and Hargreaves, 1983). Despite these descriptions, the precise timing of these structures remains poorly defined. Keon and Hargreaves (1983) described general time points in the infection process: germination and epidermal penetration by 24 hours post-inoculation (hpi), intercellular mesophyll growth by 96 hpi, and cell death by 120 hpi. Additionally, the ultrastructural events above preceded macroscopic symptoms, including water-soaked tissue, pinpoint lesions, and the eventual necrotic, netted lesions with surrounding chlorotic tissue (Keon and Hargreaves, 1983).

The heterothallic nature of *P. teres* f. *teres* has been used to develop bi-parental mapping populations, enabling the association of specific genomic loci with pathogen virulence and avirulence (Reviewed in Clare et al. 2020). The first putative effector gene locus identified in the *P. teres* f. *teres*–barley pathosystem, *AvrHar*, was mapped using the *P. teres* f. *teres* 15A × 0-1 population, where the 15A allele of *AvrHar* conferred avirulence on the barley line Harbin (Weiland et al. 1999). Two additional effector genes, *AvrPra1* and *AvrPra2*, were also mapped from the 15A × 0-1 population and conferred avirulence on the barley cultivar Prato (Lai et al. 2007). *AvrPra2* and *AvrHar* mapped to the same genomic region and were linked in repulsion (Lai et al. 2007; Wyatt et al. 2018). Another avirulence gene, Avr_Heartland_, was mapped in a *P. teres* f. *teres* population generated from two Canadian isolates that segregated for virulence on the barley cultivar Heartland (Beattie et al. 2007).

The genetics of the *P. teres* f. *teres –* barley interaction is complex, as dominant resistance, dominant susceptibility, and quantitative resistance have been identified (Reviewed in Liu et al. 2011; Clare et al. 2020). Multiple studies have mapped dominant resistance and susceptibility to *P. teres* f. *teres* to a similar genomic position on barley chromosome 6H, indicating multiple genes or alleles confer resistance or susceptibility to different pathotypes of *P. teres* f. *teres* (Reviewed in Liu et al. 2011; Clare et al. 2020). One example is the *P. teres* f. *teres* isolates 15A and 6A which show differential reactions on barley cultivars Rika and Kombar. Specifically, Rika is resistant to *P. teres* f. *teres* 15A but susceptible to 6A, whereas Kombar is resistant to 6A but susceptible to 15A (Abu Qamar et al. 2008). Genetic analysis of the Rika × Kombar doubled-haploid mapping population showed that susceptibility to each isolate was conferred by a single dominant gene (*Spt1.R* and *Spt1.K*, respectively; Abu Qamar et al. 2008; Liu et al. 2015; Richards et al. 2016). The phenotypes segregated in repulsion, indicating that the *Spt1* locus consisted of allelic variants or tightly linked genes within the same genomic region on barley chromosome 6H (Abu Qamar et al. 2008; Richards et al. 2016).

Shjerve et al. (2014) generated a bi-parental pathogen population using the *P. teres* f. *teres* isolates 15A and 6A and identified four virulence-associated quantitative trait loci (QTL) in the pathogen. 15A harbored virulence alleles at two independent QTL, *VK1* and *VK2* (<u>V</u>irulence on <u>K</u>ombar <u>1</u> and <u>2</u>), which accounted for 26% and 19% of the disease variation on Kombar, respectively. 6A harbored virulence alleles at the other two loci, *VR1* and *VR2* (<u>V</u>irulence on <u>R</u>ika <u>1</u> and <u>2</u>), which conferred virulence on Rika and accounted for 35% and 20% of the disease variation, respectively. Shjerve et al. (2014) showed that susceptibility to the *VK1*, *VK2*, and *VR2* single-virulence progeny isolates mapped to the same locus on barley chromosome 6H as the parental isolates (15A and 6A). Mapping indicated that the virulence factors underlying *VK1*, *VK2*, and *VR2*, and likely *VR1*, act as effectors by independently interacting, directly or indirectly, with the susceptibility gene(s) at the *Spt1* locus on barley chromosome 6H. Given this, we will refer to *VK1*, *VK2*, *VR1* and *VR2* as effectors.

While research continues to elucidate the genetic architecture of the *P. teres* f. *teres*-barley interaction, investigations into *in planta* colonization have stagnated since the 1980s. Although current literature provides foundational ultrastructural details, the lack of a compatible infection time-course study and the unstudied impact of specific effectors from the *P. teres* f. *teres*–barley interaction prevents a comprehensive understanding of critical infection checkpoints and the microscopic divergence between host responses. In other necrotrophic fungi, such as *Parastagonospora nodorum*, microscopic evidence has shown that necrotrophic effectors play key roles in addition to inducing programmed cell death (PCD; reviewed in Kariyawasam et al. 2023).

In the current study, we defined the *P. teres* f. *teres* infection structures and characterized the timeline of *in planta* colonization. Using this information in conjunction with *P. teres* f. *teres* genotypes that differ in virulent effector gene complements, we assessed the influence of specific effectors on overall colonization. To accomplish this, four different *P. teres* f. *teres* isolates with varying virulence profiles, based on the presence or absence of *VR1* and *VR2,* were inoculated onto the barley cultivar Rika (*Spt1.R*), which is susceptible to isolates harboring virulence alleles at the *VR1* and/or *VR2* loci. We collected samples at multiple time points and analyzed them using confocal laser scanning microscopy and machine learning-based fungal volume analysis.

## Methods

### Biological materials

The following *P. teres* f. *teres* isolates were used in this study: #5, a progeny isolate from the 15A*^vr1^*^/*vr2*^ × 6A*^VR1^*^/*VR2*^ bi-parental population that harbors the avirulent alleles at both the *vr1* and *vr2* loci (hereafter referred to as #5*^vr1/vr2^*), #60, a progeny isolate (15A*^vr1^*^/*vr2*^ × 6A*^VR1^*^/*VR2*^) that harbors the virulent allele at the *VR1* locus but the avirulent allele at the *vr2* locus (hereafter referred to as #60*^VR1/vr2^*), #72, a progeny isolate (15A*^vr1^*^/*vr2*^ × 6A*^VR1^*^/*VR2*^) that harbors the avirulent allele at the *vr1* locus but the virulent allele at the *VR2* locus (hereafter referred to as #72*^vr1/VR2^*), and 6A (syn. CA84-28-1), originally collected in Fresno County, California, USA, a parental isolate that harbors virulent alleles at both the *VR1* and *VR2* loci (hereafter referred to as 6A*^VR1/VR2^*; Wu et al. 2003; Shjerve et al. 2014; Table 1). The barley cultivar Rika (PI 269154) was used for all inoculations (Steffenson and Webster 1992).

**Table 1.** Four *P. teres* f. *teres* isolates with different complements of the effector genes *VR1* and *VR2*.

|  | **Loci alleles** |  |
|---|---|---|
| **Isolate** | *VR1* allele | *VR2* allele |
| #5 | *vr1* | *vr2* |
| #60 | *VR1* | *vr2* |
| #72 | *vr1* | *VR2* |
| 6A | *VR1* | *VR2* |

### Identification and confirmation of 15A*^vr1/vr2^* × 6A*^VR1/VR2^* progeny isolates #5*^vr1/vr2^*, #60*^VR1/vr2^*, and #72^vr1/VR2^

The *P. teres* f. *teres* 15A*^vr1/vr2^* × 6A*^VR1/VR2^* population was originally genotyped using a combination of amplified fragment length polymorphism (AFLP), simple sequence repeat (SSR), and restriction-site associated DNA –genotyping-by-sequencing (RAD-GBS) markers (Shjerve et al. 2014). The progeny isolates #5 (hereafter referred to as #5*^vr1/vr2^*), #60 (hereafter referred to as #60*^VR1/vr2^*), and #72 (hereafter referred to as #72*^vr1/VR2^*), were preliminary identified using the RAD-GBS marker set (Shjerve et al. 2014). According to the RAD-GBS marker set, #5*^vr1/vr2^*possessed the 15A*^vr1^*^/*vr2*^ alleles (avirulent) at both the *vr1* and *vr2* loci, #60*^VR1/vr2^* possessed the 6A*^VR1^*^/VR*2*^ allele (virulent) at the *VR1* locus and the 15A*^vr1/vr2^*allele (avirulent) at the *vr2* locus, and #72*^vr1/VR2^*possessed the 15A*^vr1^*^/*vr2*^ allele (avirulent) at the *vr1* locus and the 6A*^VR1^*^/VR*2*^ allele (virulent) at the *VR2* locus. Whole-genome Illumina resequencing (#5*^vr1/vr2^* NCBI Accession: SRX36034396, #72*^vr1/VR2^*, NCBI Accession: SRX36034397, and 6A*^VR11^*^/*vr2*^ NCBI Accession: SRX36034398) confirmed the genotypes at the *VR1* and *VR2* loci for each isolate. Adapter sequences were trimmed using Cutadapt 4.0 (Martin 2011). Paired reads were aligned to *P. teres* f. *teres* isolate 6A*^VR1/VR2^* (Wyatt et al. 2020) using Burrows-Wheeler Aligner (BWA) version 0.7.17 (Li 2013). The resulting sequence alignment/map (SAM) files were converted to binary alignment map (BAM) files and indexed using SAMtools (Danecek et al. 2021). We called single nucleotide polymorphisms (SNPs) using BCFtools (Danecek et al. 2021).

### Sample collection

All *P. teres* f. *teres* isolates were cultured, and inoculum was prepared as described by Koladia et al. (2017). Briefly, dried agarose plugs (stored at -20 °C) were placed on V8 PDA plates (10 g of Difco PDA, 3 g of CaCO_3_, 10 g of agar, 150 mL of V8 juice, and 850 mL of H_2_O) and incubated for five days in the dark at room temperature (RT), followed by one day under constant light at RT, and finally one day in the dark at 15 °C. Plates were flooded with sterile water and scraped with a sterile inoculation loop. Spores were counted with a hemocytometer and diluted to a concentration of 2,000 spores/mL. 60 µL of Tween 20 (J.T. Baker Chemical Co.) were added per 50 mL of spore suspension to minimize spore aggregation.

Each spore suspension was inoculated onto the leaves of the barley cultivar Rika, which had been grown in a greenhouse for 14 days or until the secondary leaf was fully expanded. Leaves were evenly inoculated with the spore suspension using a pneumatic paint sprayer (Husky) until just before runoff. Inoculated plants were placed in a mist chamber under constant light at 21 °C and 100% relative humidity for 24 hours (h). Following the incubation, plants were placed in a growth chamber at 21 °C with a 12-h photoperiod for the remainder of the experiment. We collected three leaf samples at 4, 8, 12, 24, 36, 48, 72, 96, and 120 hours post-inoculation (hpi). After each collection, we photographed the leaf tissues with a Canon EOS 70D camera mounted on a copy stand and used the samples for subsequent analysis. The experiment was conducted three times (three biological replicates) with each replicate consisting of three leaves per isolate at each timepoint.

### Confocal microscopy image analysis pipeline

The inoculated leaves outlined above were used in the confocal image analysis pipeline, as described in Nelson et al. (2024). Briefly, each leaf was cut into one 3 cm-long section and soaked in 96% ethanol until all chlorophyll pigment was removed. Once cleared, each leaf was immersed in 10% KOH at 85 °C for three minutes, then rinsed three times with 1× phosphate buffer saline (PBS) that was diluted from 10× PBS (25.6 g of Na_2_HPO_4_, 80 g of NaCl, 2 g of KCl, and 2 g of KH_2_PO_4_ per 1 liter of water, pH 7.4). Subsequently, each leaf was submerged in a 15 mL tube containing 4 mL of 1×PBS containing 0.2% (v/v) Tween20 (J.T. Baker Chemical Co.), 10 μg/mL wheat germ agglutinin labeled with fluorescein isothiocyanate (WGA-FITC, Sigma-Aldrich), and 20 μg/mL propidium iodide (PI, Sigma-Aldrich). The tubes were covered with aluminum foil and placed in a vacuum desiccator, depressurized to -100 kPa for 15 minutes using a two-stage rotary vane pump (Edwards 8.4 CDM RV12), and pressure was maintained for 3 hours. We then mounted the leaves on a microscope slide using EcoMount (Biocare Medical), cured them, and stored them at 4 °C until confocal image generation.

We analyzed the 324 mounted leaf sections using a Zeiss LSM700 laser-scanning confocal microscope (Zeiss) with objectives and laser channels as described by Nelson et al. (2024). Three images per isolate and timepoint across all biological replicates were taken using both the 20× and 40× objectives. Representative images were taken based on detailed observations collected from all leaves to minimize, but not eliminate, human bias. Confocal images (.lsm files) were imported into Imaris software v.11.0.1 (Oxford Instruments); channel emission was equalized, and images were captured from the 3D view then cropped to 102 × 102 × 46 µm (X × Y × Z) for final use. Specific z-stacks corresponding to sections inside the leaf (epidermal and mesophyll layers) were selected from the 20× magnification images using the Imaris v.11.0.1 “Delete Slices” function and used for volume analysis. *P. teres* f*. teres* hyphal volume (µm^3^) was calculated using the Imaris machine learning image-segmentation tool.

The summed hyphal volume was exported for each image and analyzed by timepoint. Image segmentation and machine-learning volume analysis can be used to interpret overarching trends in fungal volume throughout the leaf. By taking leaf sections over three biological replicates, representative confocal images were used to specifically measure how much hyphae (WGA-FITC signal) had colonized the leaf tissue. The same trained segmentation classifier was applied to all images, and cumulative statistics were analyzed and visualized to see the overall impact of *VR1* and *VR2* on *P. teres* f. *teres* hyphal volume inside the leaf. The segmentation classifier was trained on four randomly selected images from each timepoint and validated for accuracy by going through all z stacks of results and identifying inaccuracies.

### Statistical analysis

Hyphal volume data were collected for each isolate across nine time points (4, 8, 12, 24, 36, 48, 72, 96, and 120 hpi) and used as a proxy for cumulative fungal volume (µm^3^). Total measurements were compiled into an Excel spreadsheet containing columns for isolate, replicate, time (hpi), and volume (µm³) then uploaded into JMP Student Edition 19 (JMP Statistical Discovery LLC). Isolate and replicate were designated as categorical factors, volume (µm³) as a continuous response variable, and the data were sorted by time. Data were analyzed separately for each timepoint using time as a “By” variable. A general linear model was fitted using the Standard Least Squares method within JMP’s “Fit Model” platform. We calculated least-squares means (LSMeans) and their standard errors and performed pairwise comparisons to identify significant differences among all isolate combinations. If the analysis of variance (ANOVA) was significant (alpha=0.05), we conducted isolate effect tests and post-hoc comparisons using Tukey’s honestly significant difference (HSD) test. LSMeans plots with significance following Tukey’s HSD were generated to visualize the data.

## Results

### *P. teres* f. *teres* infection structure signatures

Confocal analysis identified specific structures of *P*. *teres* f. *teres* infection on barley. The infection structures included formation of an appressoria-like structure, penetration into the epidermal cell, formation of an intracellular vesicle and appendages, penetration into the mesophyll apoplastic space, intercellular hyphal growth in the mesophyll, rapid hyphal growth parallel and perpendicular to the leaf length, hyphal stratification, and pericellular encasement (Figure 1).

**Figure 1:**
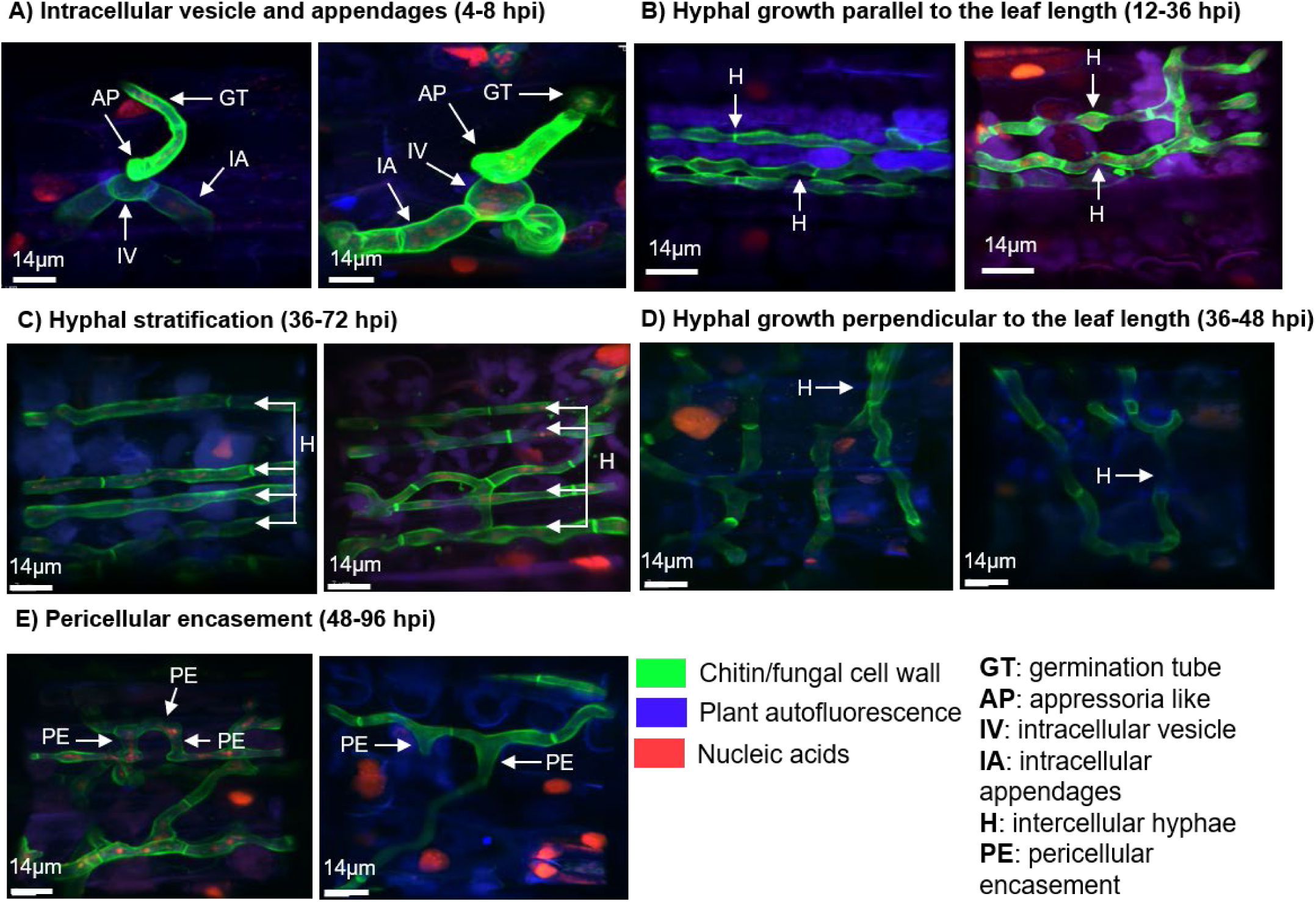
*Pyrenophora teres* f. *teres* growth development signatures. **A.** From 4-8 hpi, an intracellular vesicle (IV) forms inside the epidermal cell under the appressoria-like (AP) penetration point. From the vesicle, intracellular appendages (IA) grow and penetrate the mesophyll layer by 12 hpi. **B.** Intercellular hyphae grow from the intracellular appendages (IA), first extending vertically, then parallel to the leaf length in the mesophyll layer from 12 to 36 hpi. **C.** From 36 to 72 hpi hyphal stratification occurs where three or more parallel intercellular hyphae (H) aggregate within a single mesophyll layer. **D.** Between 36 and 48 hpi, intercellular hyphae (H) grow perpendicular to the leaf length, extending three or more mesophyll cells. **E.** Pericellular encasement (PE) occurs from 48 to 96 hpi, where hyphae (H) converge, circumferentially ensheathe mesophyll cells, and expand into the available apoplastic space.

By 8 hours post-inoculation (hpi), an appressoria-like structure (AP) emerged from a germinated conidium on the leaf surface, then penetrated the epidermal cell (Figure 1A). Inside the epidermal cell, the formation of an intracellular vesicle (IV) and intracellular appendages had already occurred (IA; Figure 1A). The intracellular vesicle (IV) consisted of a central, bulbous structure with two to three thick, leg-like intracellular appendages protruding from it (IA; Figure 1A). Inside the intracellular vesicle, numerous small propidium iodide stained (red) dots, consistent with nuclei, were distributed throughout the structure (Figure 1A). By 12 hpi, the intracellular appendages (IA) penetrated the mesophyll cell layer where hyphae grew vertically and intercellularly (Figure 1A-B). By 36 hpi, the intercellular hyphae grew at a rapid rate parallel to the leaf length, spanning multiple mesophyll cells (Figure 1B). By 48 hpi, hyphal stratification, which consisted of three or more independent hyphae growing parallel between mesophyll cells in the same concentrated area, was observed (Figure 1C). Additionally, hyphal growth perpendicular to the leaf length was also observed and resulted in an increasingly dense grid-like colonization of the mesophyll layer (Figure 1D). By 72 hpi, hyphae exhibited pericellular encasement, where they expanded into the available apoplastic space between mesophyll cells and fully surrounded them (Figure 1E).

### Comparative *P. teres* f. *teres* infection timeline

Having established *P. teres* f. *teres’* specific infection structures and their timelines, we next compared how isolates with different virulence profiles progressed through them. Macro- and microscopic fungal activity along with host responses were analyzed at 4, 8, 12, 24, 36, 48, 72, 96, and 120 hours post-inoculation (hpi) using the validated *P. teres* f. *teres* isolates #5*^vr1/vr2^*, #60*^VR1/vr2^*, #72*^vr1/VR2^*, and 6A*^VR1^*^/*VR2*^ (Shjerve et al. 2014; Figures 2-5). Whole-genome Illumina resequencing confirmed that each isolate harbored the expected *VR1* and *VR2* alleles at each locus. #5*^vr1/vr2^*possesses the 15A*^vr1^*^/*vr2*^ allele (avirulent) at the *vr1* locus and the 15A*^vr1^*^/*vr2*^ allele (avirulent) at the *vr2* locus; #60*^VR1/vr2^* possesses the 6A*^VR1^*^/VR*2*^ allele (virulent) at the *VR1* locus and the 15A*^vr1/vr2^*allele (avirulent) at the *vr2* locus; and #72*^vr1/VR2^* possesses the 15A*^vr1^*^/*vr2*^ allele (avirulent) at the *vr1* locus and the 6A*^VR1^*^/VR*2*^ allele (virulent) at the *VR2* locus. We chose these isolates to compare how the effector genes *VR1* and *VR2* influenced the timing and severity of host colonization and lesion development.

**Figure 2:**
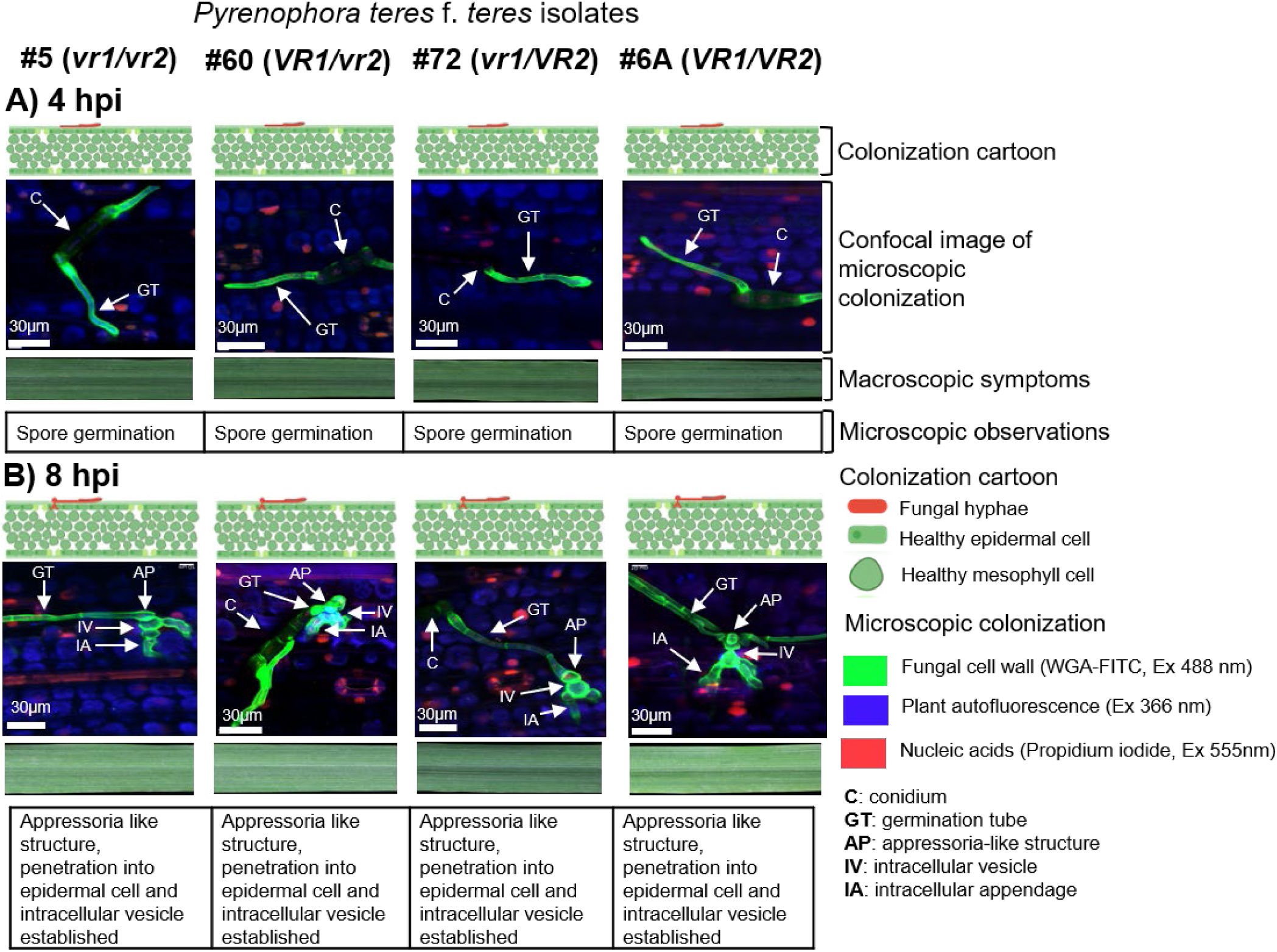
Disease progression of *Pyrenophora teres* f. *teres* isolates #5*^vr1/vr2^*, #60*^VR1/vr2^*, #72*^vr1/VR2^*, and 6A*^VR1/VR2^* on barley cultivar Rika 4 and 8 hours post inoculation (hpi). Macroscopic and microscopic disease progression on the susceptible barley line Rika was analyzed and compared among four isolates at 4 and 8 hpi. **A.** At 4 hpi all four isolates had a conidium (C) that initiated germination via a germination tube (GT) and no macroscopic leaf symptoms were visible. **B.** At 8 hpi, all isolates had germination tubes (GT) that differentiated into an appressoria-like structure (AP), then penetrated an epidermal cell, where an intracellular vesicle (IV) and intracellular appendages (IA) were established. Macroscopic symptoms remained absent from all leaves. Cartoons created in BioRender. Nelson, A. (2027) https://BioRender.com/km16xfb

#### 4 to 24 hours post inoculation (hpi)

Colonization patterns between 4 and 24 hpi were similar across all isolates (Figures 2 and 3). By 4 hpi, germination tubes emerged from the conidium (Figure 2A). Between 4 and 8 hpi, an appressoria-like structure formed at the end of the germination tube, penetrated the epidermal cell, and established an intracellular vesicle with two to three appendages inside the cell (Figure 2B). Between 8 and 12 hpi, the intracellular appendages penetrated the mesophyll layer (Figure 3A). By 24 hpi, intercellular hyphae had grown vertically and began growth parallel to the leaf length (Figure 3B).

**Figure 3:**
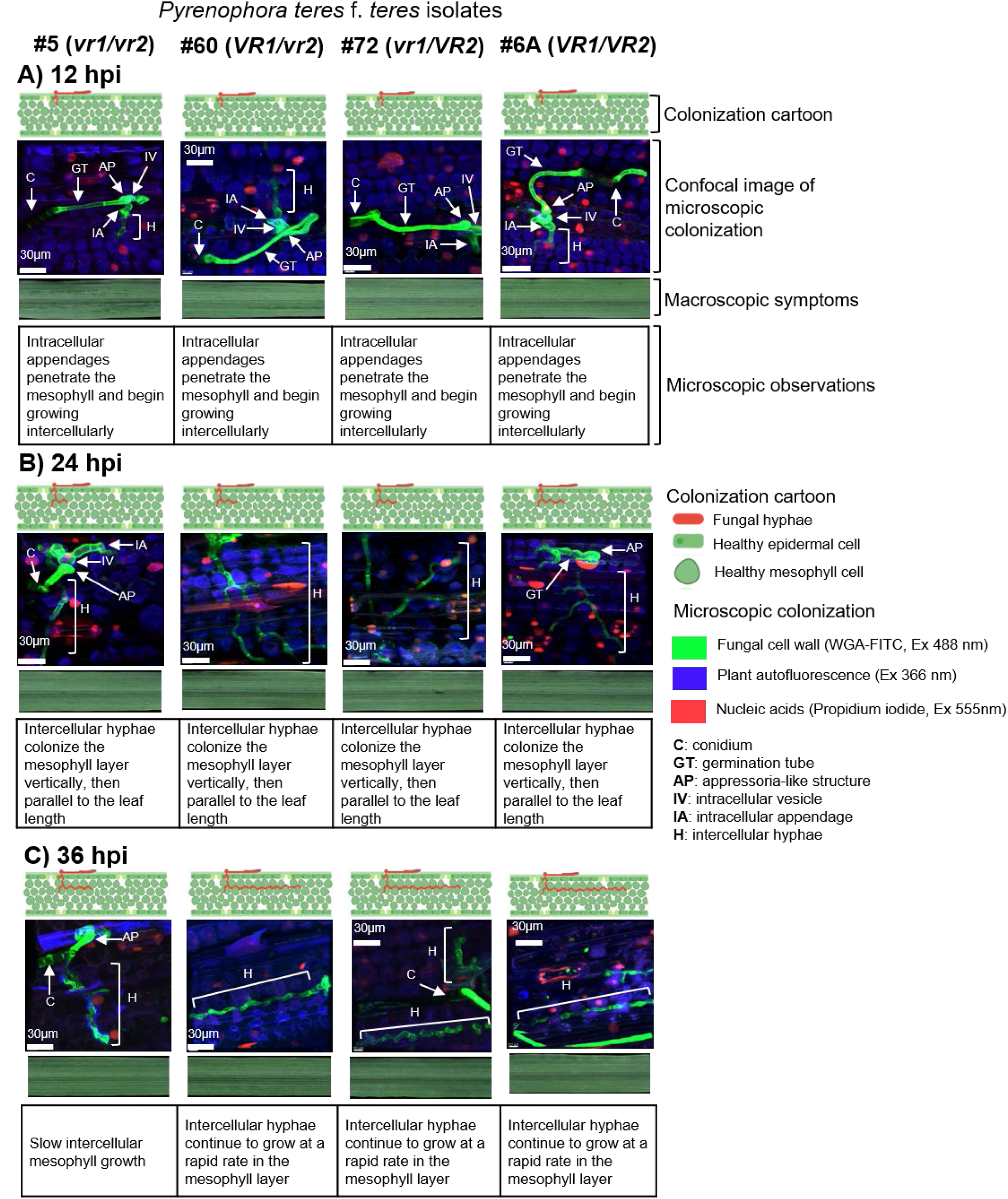
Disease progression of *Pyrenophora teres* f. *teres* isolates #5*^vr1/vr2^*, #60*^VR1/vr2^*, #72*^vr1/VR2^*, and 6A*^VR1/VR2^* on barley cultivar Rika 12 to 36 hours post inoculation (hpi). Macroscopic and microscopic disease progression on the susceptible barley line Rika was analyzed and compared among four isolates at 12-36 hpi. **A.** At 12 hpi, intracellular appendages (IA) extend from the intracellular vesicle (IV) into the mesophyll layer, initiating intercellular hyphae (H) across all isolates. Macroscopic lesions are absent. **B.** At 24 hpi, hyphae (H) continue to colonize the mesophyll layer, expanding parallel to the leaf length across all isolates and no macroscopic lesions are visible. **C.** At 36 hpi, in isolates #60*^VR1/vr2^*, #72*^vr1/VR2^*, and 6A*^VR1/VR2^*, hyphae (H) continue to colonize the mesophyll, but in isolate #5*^vr1/vr2^* growth is noticeably slower, and macroscopic lesions remain absent across all isolates. Cartoons created in BioRender. Nelson, A. (2027) https://BioRender.com/km16xfb.

#### 36 to 72 hours post inoculation (hpi)

By 36 hpi, in leaves inoculated with #60*^VR1/vr2^*, #72*^vr1/VR2^*, and 6A*^VR1/VR2^* intercellular hyphae had rapidly grown parallel to the leaf length (Figure 3C). Leaves inoculated with #5*^vr1/vr2^* also grew parallel to the leaf length but at a slower rate (Figure 3C). We also captured macroscopic images of Rika barley inoculated with each isolate at all time points, then evaluated them for leaf lesion development. No visible lesion development was observed on any leaves by 36 hpi (Figures 2 and 3).

Between 36 and 48 hpi, differences between the avirulent #5*^vr1/vr2^*and the virulent #60*^VR1/vr2^*, #72*^vr1/VR2^*, and 6A*^VR1/VR2^* became pronounced (Figure 4). In leaves inoculated with the virulent isolates, hyphae had grown rapidly in the mesophyll layer parallel to the leaf length, along with hyphal stratification and the initiation of perpendicular growth from the parallel hyphae. #5*^vr1/vr2^*showed both parallel and perpendicular growth in the mesophyll layer but was less extensive relative to the virulent isolates (Figure 4A; Nelson et al. 2024). In leaves inoculated with the virulent isolates (#60*^VR1/vr2^*, #72*^vr1/VR2^*, and 6A*^VR1/VR2^*), nuclear breakdown, indicating PCD, was observed (Figure 4A; Danon et al. 2000; Domínguez and Cejudo, 2015). Nuclear breakdown was determined by the appearance of fragmented red (propidium iodide) throughout the mesophyll cell, which resulted in a vibrant purple color (Figures 4 and 5). However, #5*^vr1/vr2^*showed no cytological features associated with PCD (Figure 4A). Leaves inoculated with #5*^vr1/vr2^* showed no macroscopic symptoms (Figure 4A). However, leaves inoculated with #60*^VR1/vr2^* and #72*^vr1/VR2^* showed the first signs of lesion development through small chlorotic spots (Figure 4A). Leaves inoculated with 6A*^VR1/VR2^* developed more pronounced chlorotic and necrotic spots (Figure 4A).

**Figure 4:**
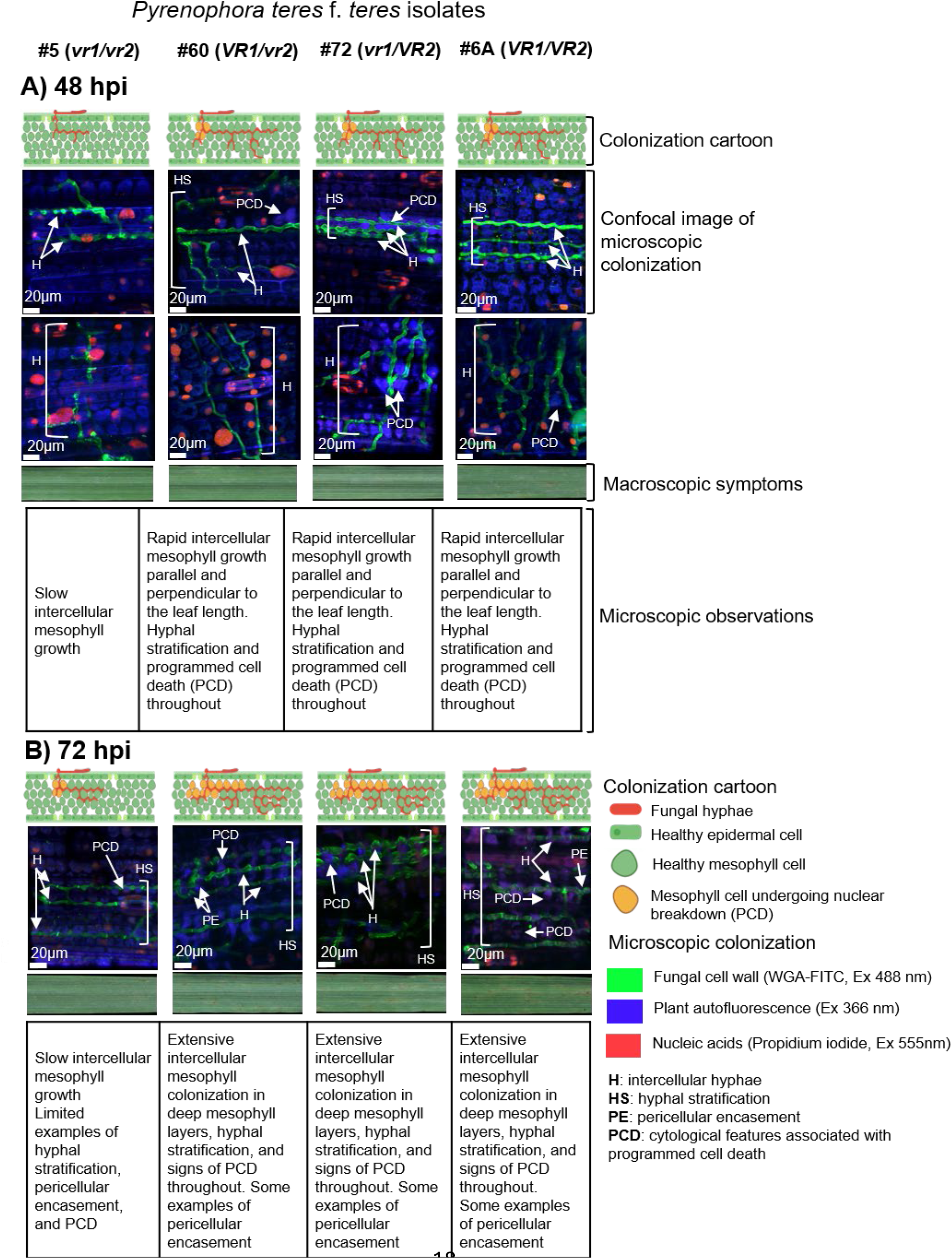
Disease progression of *Pyrenophora teres* f. *teres* isolates #5*^vr1/vr2^*, #60*^VR1/vr2^*, #72*^vr1/VR2^*, and 6A*^VR1/VR2^* on barley cultivar Rika 48 and 72 hours post inoculation (hpi). Macroscopic and microscopic disease progression on the susceptible barley line Rika was analyzed and compared among four isolates at 48 and 72 hpi. **A.** At 48 hpi, the top image depicts intercellular hyphal growth (H) parallel to the leaf length and the bottom image depicts intercellular hyphal growth (H) perpendicular to the leaf length. Isolates #60*^VR1/vr2^*, #72*^vr1/VR2^*, and 6A*^VR1/VR2^* had rapid parallel and perpendicular hyphal growth (H), hyphal stratification (HS), signs of PCD, and small, circular lesions on the leaves. In contrast, isolate #5*^vr1/vr2^* maintained slow hyphal growth (H) and no signs of PCD or macroscopic lesions. **B.** At 72 hpi, isolates #60*^VR1/vr2^*, #72*^vr1/VR2^*, and 6A*^VR1/VR2^* continued extensive growth with hyphal stratification (HS), PCD, and expanded oval lesions with light chlorosis. Isolate #5*^vr1/vr2^* maintained slow hyphal growth (H) with isolated signs of PCD, and leaves developed small chlorotic flecks. Cartoons created in BioRender. Nelson, A. (2027) https://BioRender.com/km16xfb.

By 72 hpi, mesophyll colonization was more extensive in leaves inoculated with #60*^VR1/vr2^*, #72*^vr1/VR2^*, and 6A*^VR1/VR2^*, including extensive parallel and perpendicular growth into deep mesophyll layers, hyphal stratification, pericellular encasement, and PCD (Figure 4B). In leaves inoculated with #5*^vr1/vr2^*, mesophyll colonization continued to lag, although limited examples of hyphal stratification and pericellular encasement were still observed (Figure 4B). Leaves inoculated with #60*^VR1/vr2^*, #72*^vr1/VR2^*, and 6A*^VR1/VR2^* developed oval necrotic lesions surrounded by chlorosis (Figure 4B). Additionally, necrotic lesions in leaves inoculated with, #72*^vr1/VR2^* and 6A*^VR1/VR2^*began to expand longitudinally (Figure 4B). Leaves inoculated with #5*^vr1/vr2^*developed small chlorotic flecks (Figure 4B).

#### 96 to 120 hours post inoculation (hpi)

By 96 hpi, leaves inoculated with #60*^VR1/vr2^* and #72*^vr1/VR2^* continued to show mesophyll colonization throughout all layers, with hyphal stratification, pericellular encasement, PCD, and collapsed mesophyll cells (Figure 5A). Leaves inoculated with 6A*^VR1/VR2^*showed widespread hyphal colonization in deep mesophyll layers, including hyphal stratification, pericellular encasement, PCD, and collapsed mesophyll cells (Figure 5A). Leaves inoculated with #5*^vr1/vr2^* had slow mesophyll colonization with limited examples of hyphal stratification, pericellular encasement, and PCD (Figure 5A; Danon et al. 2000; Domínguez and Cejudo, 2015). The above indicates that VR1 and VR2 are involved in facilitating colonization and inducing cell death.

**Figure 5:**
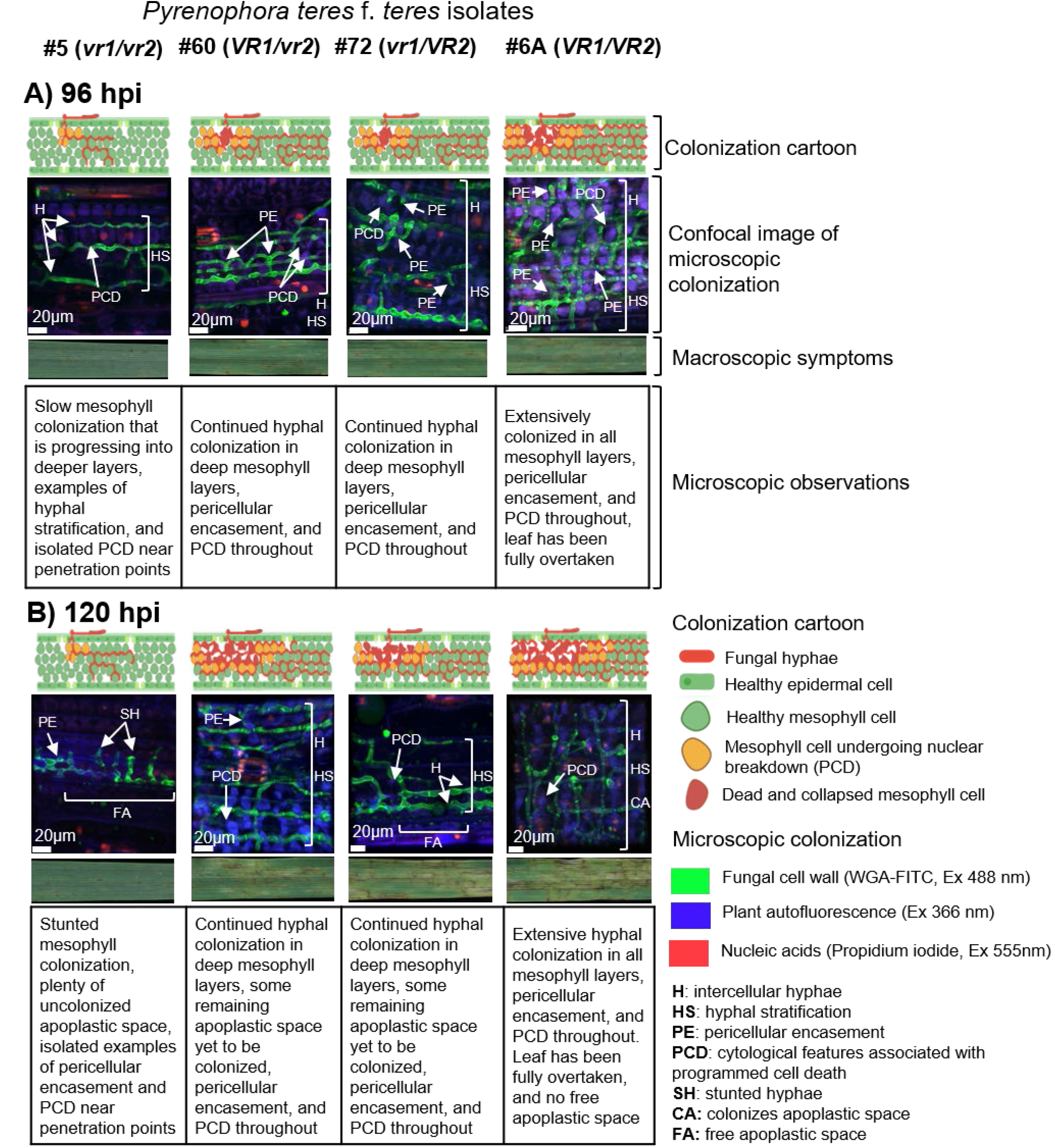
Disease Progression of *Pyrenophora teres* f. *teres* isolates #5*^vr1/vr2^*, #60*^VR1/vr2^*, #72*^vr1/VR2^*, and 6A*^VR1/VR2^* on barley cultivar Rika at 96 and 120 hours post inoculation (hpi). Macroscopic and microscopic disease progression on the susceptible barley line Rika was analyzed and compared among four isolates at 96 and 120 hpi **A.** At 96 hpi, isolates #60*^VR1/vr2^*, #72*^vr1/VR2^*, and 6A*^VR1/VR2^*had extensive hyphal colonization (H, HS), pericellular encasement (PE) and PCD culminating in net-like lesions with surrounding chlorosis. Isolate 6A*^VR1/VR2^*had higher hyphal abundance than isolates #60*^VR1/vr2^* and #72*^vr1/VR2^*. Isolate #5*^vr1/vr2^* exhibited reduced hyphal colonization (H), isolated PCD, and leaves had small circular necrotic lesions. **B.** At 120 hpi, isolates #60*^VR1/vr2^*, #72*^vr1/VR2^*, and 6A*^VR1/VR2^* continued extensive hyphal colonization (H, HS) and PCD. Isolate 6A*^VR1/VR2^* exhibited the most complete apoplastic colonization (CA) and extensive net-like lesions while isolates #60*^VR1/vr2^* and #72*^vr1/VR2^* had more uncolonized apoplastic space (FA) and isolate #60*^VR1/vr2^* had slightly more hyphae than isolate #72*^vr1/VR2^*. Isolate #*5^vr1/vr2^* exhibited stunted hyphae (SH) and small circular necrotic lesions. Cartoons created in BioRender. Nelson, A. (2027) https://BioRender.com/km16xfb.

The 96 hpi *in planta* colonization patterns were mirrored at the macroscopic level. Leaves inoculated with #60*^VR1/vr2^* and #72*^vr1/VR2^* exhibited expanding necrotic lesions with surrounding chlorosis that began to develop the characteristic netting morphology (Figure 5A). In leaves inoculated with 6A*^VR1/VR2^*, necrotic lesions coalesced and developed a net-like appearance surrounded by chlorosis throughout the leaf (Figure 5A). Meanwhile, only small circular necrotic lesions were observed on leaves inoculated with #5*^vr1/vr2^* (Figure 5A).

Between 96 and 120 hpi, colonization patterns persisted. Leaves inoculated with #60*^VR1/vr2^* and #72*^vr1/VR2^* continued to expand their extensive colonization of the free apoplastic space into the mesophyll layer (Figure 5B). However, some areas remained uncolonized, and PCD was less extensive relative to 6A*^VR1/VR2^* (Figure 5B). Leaves inoculated with #60*^VR1/vr2^*had colonized more mesophyll relative to leaves inoculated with #72*^vr1/VR2^*(Figure 5B). In leaves inoculated with 6A*^VR1/VR2^*, hyphae formed an extensive grid-like pattern and colonized most of the apoplastic space in the mesophyll layer (Figure 5B). Pericellular encasement and PCD were present throughout, and hyphae had overtaken and necrotized the barley leaf (Figure 5B). Leaves inoculated with #5*^vr1^*^/*vr2*^ continued to slowly colonize the mesophyll, with limited examples of pericellular encasement and PCD (Figure 5B).

When evaluating lesion development at the 120 hpi timepoint, leaves inoculated with #60*^VR1^*^/*vr2*^ and #72*^vr1^*^/*VR2*^ developed coalescing necrotic lesions that progressed transversely and longitudinally, with surrounding chlorotic areas, displaying classic net-like necrotic lesions (Figure 5B). Leaves inoculated with 6A*^VR1^*^/*VR2*^ showed more extensive damage with most of the leaf overtaken by necrotic, net-like lesions with accompanying chlorosis (Figure 5B). Conversely, leaves inoculated with #5*^vr1^*^/*vr2*^ continued to show only a few small circular necrotic lesions (Figure 5B).

Together the above observations suggest that *VR1* and *VR2* each promote colonization and that their combination produces the most rapid and extensive colonization. The independent effect is highlighted by the microscopic colonization differences and macroscopic lesion formation between #5*^vr1/vr2^*, #60*^VR1/vr2^* and #72*^vr1/VR2^*. The combined effect is shown by the more rapid and extensive colonization of 6A*^VR1/VR2^*compared with both #60*^VR1/vr2^* and #72*^vr1/VR2^*.

### Machine-learning-based volume analysis

To quantify fungal colonization, image segmentation machine-learning-based volume analysis was performed for each isolate (#5*^vr1/vr2^*, #60*^VR1/vr2^*, #72*^vr1/VR2^*, and 6A*^VR1/VR2^*) across nine time points (4, 8, 12, 24, 36, 48, 72, 96, and 120 hpi; Figure 6). We analyzed the summed volume from three representative images per isolate-timepoint combination to determine average hyphal volume (µm^3^). Significant ANOVAs with isolate and replicate as factors revealed that hyphal volume varied significantly among isolates at 72 (*p* <0.05), 96 (*p* <0.05), and 120 hpi (*p* <0.01). Effect tests were used in conjunction with significant ANOVAs to isolate and evaluate the contribution of each main effect and interaction term. The effect tests were significant for the isolate term at 72 (*p* <0.05), 96 (*p* <0.05), and 120 hpi (*p* <0.01). Consequently, we used a post-hoc Tukey’s honestly significant difference (HSD) test to compare fungal volume and the relative impact of VR1 and VR2 across isolates (Tables A1, A2, and A3).

**Figure 6:**
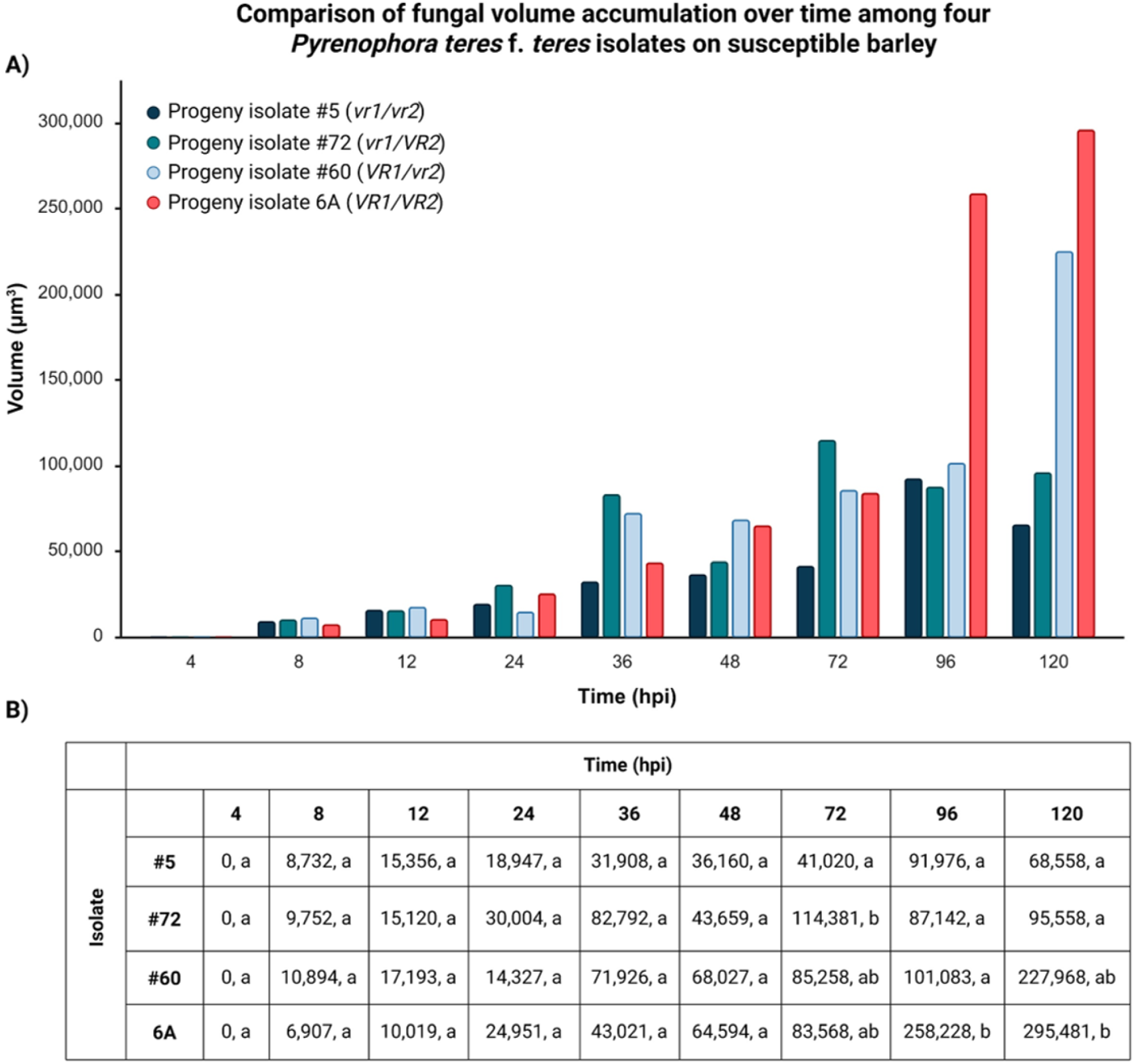
Fungal volume accumulation among four *Pyrenophora teres* f. *teres* isolates on susceptible barley. **A**. Fungal volume (*y*-axis) was quantified for the 15A×6A progeny isolates #5*^vr1^*^/*vr2*^, #72*^vr1^*^/*VR2*^ and #60*^VR1^*^/*vr2*^ and the parental isolate 6A*^VR1^*^/*VR2*^, from 4 to 120 hours post inoculation (hpi; *x-*axis) on the susceptible barley cultivar Rika. Column colors represent individual isolates; navy blue (#5*^vr1^*^/*vr2*^), teal (#72*^vr1^*^/*VR2*^), light blue (#60*^VR1^*^/*vr2*^), salmon (6A*^VR1^*^/*VR2*^). **B.** The LSMeans for each isolate (rows) across all timepoints (columns). Letters indicate significant differences (*p* <0.05) within each timepoint based on analysis of variance using a general linear model followed by a Tukey’s honestly significant differences (HSD) test. Each timepoint was analyzed independently and statistical significance cannot be compared between timepoints. Created in BioRender. Nelson, A. (2027) https://BioRender.com/24apmaw.

Hyphal volume did not significantly differ among isolates from 4 to 48 hpi (Figure 6), although microscopic and macroscopic differences were apparent at 36 and 48 hpi (Figure 3C, Figure 4A). At 72 hpi, #72*^vr1/VR2^* had significantly more hyphal volume relative to #5*^vr1/vr2^*, but there was no significant difference in hyphal volume compared to #60*^VR1/vr2^* and 6A*^VR1/VR2^*(Figure 6; Table A1). At 96 hpi, #5*^vr1/vr2^*, #72*^vr1/VR2^*, and #60*^VR1/vr2^* were not significantly different from one another but had significantly lower hyphal volume relative to 6A*^VR1/VR2^* (*p* <0.05; Figure 6; Table A2). At 120 hpi, #5*^vr1/vr2^* and #72*^vr1/VR2^*continued to have significantly lower hyphal volume (*p* <0.01) relative to 6A*^VR1/VR2^* (Figure 6; Table A3).

Although hyphal volume for #5*^vr1/vr2^*, #72*^vr1/VR2^*, and #60*^VR1/vr2^* was not significantly different from one another at 96 hpi and 120 hpi, #60*^VR1/vr2^* had hyphal volume that was trending higher relative to #5*^vr1/vr2^* and #72*^vr1/VR2^*but lower than 6A*^VR1/VR2^* (Figure 6; Table A3). At 120 hpi, 6A*^VR1/VR2^*had the largest mean hyphal volume with significantly more hyphal volume than #5*^vr1/vr2^* and #72*^vr1/VR2^* (*p* <0.01; Figure 6; Table A3). These data suggest a correlation between the presence of *VR1* and *VR2* and *in planta* fungal fitness.

Isolates that had the virulent *VR1* allele (#60*^VR1/vr2^* and 6A*^VR1/VR2^*) had larger hyphal volume LSMeans at later timepoints compared to isolates that either had no virulence alleles or only the virulent *VR2* allele (#5*^vr1/vr2^* and #72*^vr1/VR2^*, Figure 6). This indicates that *VR1* has a larger effect on colonization relative to *VR2* at later timepoints (Figure 6), which aligns with the higher association with disease for *VR1* (35%) compared to *VR2* (20%), as reported by Shjerve et al. (2014). When both *VR1* and *VR2* virulence alleles were present, as in 6A*^VR1/VR2^*, there was a significant increase in hyphal volume compared to #5*^vr1/vr2^* and #72*^vr1/VR2^*at later timepoints (*p* <0.05; Figure 6; Tables A2-3). These results suggest that *VR1* alone plays a major role in *P. teres* f*. teres* colonization at later timepoints, but when both *VR1* and *VR2* are present, they act synergistically to maximize colonization (Figure 6).

## Discussion

Many studies have explored the complex genetics of the *P. teres* f. *teres* – barley interaction, revealing the involvement of dominant resistance, dominant susceptibility, and quantitative resistance in the host (Reviewed in Liu et al. 2011; Clare et al. 2020). Previous microscopic ultrastructural analyses (Keeling and Banttari, 1975; Van Caeseele and Grumbles, 1979; Keon and Hargreaves, 1983) established the foundational framework for understanding the *P. teres* f. *teres* infection process. However, these earlier analyses did not define the timing of individual infection structures, a critical infection timeline, or the impact of *P. teres* f. *teres* effectors on colonization. To address these gaps, we first defined the infection structures and compatible infection timeline of the *P. teres* f. *teres*–barley pathosystem using confocal laser scanning microscopy (CLSM). Second, we compared four *P*. *teres* f. *teres* isolates with different effector genes using macroscopic lesion development, microscopic *in planta* colonization, and fungal volume quantification across multiple timepoints.

### *Pyrenophora teres* f. *teres* infection structure signatures

CLSM analyses revealed specific infection structures across all *P. teres* f. *teres* isolates. These structures included appressoria-like structures, intracellular vesicles, intracellular appendages, intercellular mesophyll growth (both parallel and perpendicular to the leaf length), hyphal stratification, and pericellular encasement (Figure 1). Many of these structures corroborate observations from previous electron microscopy studies (Keeling and Banttari, 1975; Van Caeseele and Grumbles, 1979; Keon and Hargreaves, 1983), however, we identified some nuances.

#### Germination and penetration into epidermal cells

Conidia contained two to three septa, with nucleic material inside (stained by propidium iodide in red; Figure 2A). Germination tubes extended from the conidium, followed by concentrated areas of nucleic material throughout the developing germination tube (Figure 2A; Van Caeseele and Grumbles, 1979; Keon and Hargreaves, 1983). Germination tubes extended across the leaf surface, developing a club-shaped appressoria-like structure above the epidermal layer, and subsequently penetrated the epidermal cell (Figure 1A; Figure 2B). We considered the structure appressoria-like because a penetration peg was not visible and a penetration peg is a critical component of a true appressorium (Perez-Nadales et al. 2014; Figures 1A and 2B). However, CLSM has lower resolution, and a fine penetration peg could be missed. Earlier studies by Van Caeseele and Grumbles (1979) and Keon and Hargreaves (1983) identified this structure as an appressorium and reported a penetrating hypha responsible for breaching the cuticle and the underlying epidermal cell wall. However, capturing penetration hyphae with electron microscopy can be difficult because of the thin cell wall and interference from embedding resin, making structural differentiation challenging (Mendgen and Deising, 1993). The definitive presence of a penetration peg needs to be determined using higher resolution microscopy to confirm the findings of previous studies (Van Caeseele and Grumbles, 1979; Keon and Hargreaves, 1983)

Using previous electron microscopy studies, we can fill in some of the ultrastructural gaps from our CLSM infection timeline. During epidermal cell penetration, a round, smooth pore with an amorphous layer and Woronin bodies was established at the host cuticle interface (Van Caeseele and Grumbles, 1979; Keon and Hargreaves, 1983). Keon and Hargreaves (1983) also documented that upon penetration, the free edges of the plant cuticle were invaginated into the epidermal cell wall, with occasional detachment from the underlying cell wall. Concurrently, the walls of penetrated epidermal cells and parts of adjacent epidermal cell walls became impregnated with electron-dense material; notably, no electron-dense material was found directly around the penetration sites (Van Caeseele and Grumbles, 1979; Keon and Hargreaves, 1983).

#### Intracellular vesicle and appendages

Following penetration of the epidermal cell, an intracellular vesicle formed, with electron-dense material accumulating at its neck (Figure 1A; Figure 2B; Van Caeseele and Grumbles, 1979; Keon and Hargreaves, 1983). The intracellular vesicle appeared as a single, bulbous structure (Figure 1A; Figure 2B), enveloped by an electron-dense amorphous layer containing nucleic acid material and dense cytoplasm rich in organelles (Van Caeseele and Grumbles, 1979; Keon and Hargreaves, 1983). However, a large central vacuole subsequently developed, leading to the apparent loss of cytoplasm and organelles (Van Caeseele and Grumbles, 1979; Keon and Hargreaves, 1983). While Van Caeseele and Grumbles (1979) termed this “infection vesicle,” we have adopted the “intracellular vesicle” terminology, aligning with Keon and Hargreaves (1983), as it better describes the observed growth characteristics. The formation of the intracellular vesicle within the epidermal cell led to a complete loss of membrane integrity in the epidermal cell, and adjacent cells exhibited cytoplasmic vesiculation (Van Caeseele and Grumbles, 1979; Keon and Hargreaves, 1983).

*P. teres* f. *teres’* intracellular vesicle has structural similarities to substomatal vesicles in *Puccinia* species, and intracellular/infection hyphae from *Colletotrichum* and *Magnaporthe* species (Perfect and Green 2001). Intracellular/infection hyphae have been implicated in suppressing plant defense responses and nutrient uptake via ATPases and sugar gradients (Spencer-Phillips and Gay, 1981; Heath and Skalamera, 1997; Perfect and Green, 2001). Given the structural and location similarities, we propose that *P. teres* f. *teres’* intracellular vesicle may function as a feeding structure, facilitating the initial transfer of nutrients. Additionally, we agree with Keon and Hargreaves (1983) that it may also help overcome host defenses. Notably, Keon and Hargreaves (1983) reported a second, larger intracellular vesicle; however, Van Caeseele and Grumbles (1979) did not observe any secondary vesicle formation. We observed secondary intracellular vesicle formation, but it was rare and inconsistent. It is still unclear whether secondary intracellular vesicles develop under specific conditions during *P. teres* f. *teres* infection or if they simply occur at random.

From the intracellular vesicle, two to three intracellular appendages emerged, establishing themselves inside the epidermal cell before penetrating the mesophyll layer and forming a septum at the epidermal cell exit site (Figure 1A; Figure 2B; Van Caeseele and Grumbles, 1979; Keon and Hargreaves, 1983). These appendages were notably thicker than hyphae observed in the mesophyll layer (Figures 1A and 2B; Keon and Hargreaves, 1983). While Keon and Hargreaves (1983) termed these structures “infection hyphae” because they penetrated the mesophyll layer, we updated the term to “intracellular appendages” to better reflect their distinct, thicker morphology and direct attachment to the intracellular vesicle.

#### Mesophyll colonization

Upon penetration of the mesophyll layer, the intercellular hyphae initially grew vertically (Figures 3A and 3B) before establishing growth parallel to the leaf length (Figure 1B; Figure 3C; Van Caeseele and Grumbles, 1979; Keon and Hargreaves, 1983). While Van Caeseele and Grumbles (1979) reported hyphal growth through mesophyll cells prior to intercellular establishment, we interpret their observation as the initial vertical growth phase. In our observations, intercellular mesophyll growth conformed to the shape of the mesophyll cells, exhibiting thinning to navigate between cells and thickening to occupy the free apoplastic space (Figure 1B; Figure 3C). We observed multiple layers of parallel intercellular hyphal growth, which we termed “hyphal stratification” (Figure 1C; Figure 4A and B). Soon after the establishment of hyphal stratification, we observed intercellular hyphal growth perpendicular to the leaf length. Perpendicular growth appears to establish new areas of colonization that later undergo PCD, contributing to the netted lesion characteristic of *P. teres* f. *teres* (Figure 1D; Figure 4A).

Eventually, multiple intercellular hyphae converged around individual mesophyll cells, increasing hyphal surface area and ultimately surrounding the cell (Figure 1E; Figure 5A and B). We designated this growth type “pericellular encasement” where “pericellular” describes the targeting and surrounding of individual mesophyll cells, while encasement reflects how hyphae adapt their shape to surround the cell (Figure 1E; Figure 5A and B). Keon and Hargreaves (1983) noted that intercellular hyphae were attached to host cell walls by a deposit of electron-lucent material, which may explain how *P. teres* f. *teres* molds itself around mesophyll cells. The electron-lucent material could be a fungal extracellular matrix of polysaccharides, glycoproteins, and glucan, which has been shown to act as a biological glue that attaches fungal hyphae to plant cells (Heller 2020). The close contact between hyphae and plant cells may position the *P. teres* f. *teres* to rapidly absorb nutrients once host cell death is induced. Nutrient uptake directly fuels pathogen reproduction via sporulation, facilitating spread to new leaves and perpetuating the polycyclic disease cycle.

#### Cytological features associated with programmed cell death (PCD)

Mesophyll colonization resulted in the appearance of cytological features associated with PCD (Figure 4; Danon et al. 2000; Domínguez and Cejudo, 2015). Consistent with Keon and Hargreaves (1983), our observations revealed nuclear breakdown, which is indicative of PCD (Figure 4; Danon et al. 2000; Domínguez and Cejudo, 2015). Keon and Hargreaves (1983) further described ultrastructural signs of PCD that were beyond the resolution of CLSM analysis. Observations included vesiculation (the formation of vesicles within cells that are used to remove damaged membrane patches), membrane disruption, a disorganized cytoplasm with dense osmophilic bodies, increased numbers of mitochondria, vesiculated endoplasmic reticulum, and electron-dense bodies surrounding cells. Furthermore, consistent with previous studies (Van Caeseele and Grumbles, 1979; Keon and Hargreaves, 1983), mesophyll cells exhibiting cytological features associated with PCD and/or cell collapse were in direct contact with intercellular hyphae, many of which showed hyphal stratification and pericellular encasement (Figure 5). Additionally, cytological features associated with PCD and/or cell collapse consistently appeared behind advancing hyphae, supporting the argument that necrotrophic pathogens can strategically manipulate PCD induction rather than indiscriminately releasing cell wall-degrading enzymes or effectors during *in planta* colonization (Kariyawasam et al. 2022).

#### Compatible *P. teres* f. *teres* infection model

While our observations largely aligned with the microscopic studies of *P. teres* f. *teres* including Keeling and Banttari (1975), Van Caeseele and Grumbles (1979), and Keon and Hargreaves (1983), one study reported markedly different findings. Lightfoot and Able (2010) stated that susceptible barley plants inoculated with *P. teres* f. *teres* did not establish any intracellular growth in the epidermal layer; instead, it grew sub-epidermally or directly into the mesophyll layer to establish intercellular growth. However, our analysis of 324 leaf samples inoculated with four distinct *P. teres* f. *teres* isolates revealed no instances of bypassing intracellular epidermal growth and always included the formation of the intracellular vesicle and appendages (Figure 1A; Figure 2B). Furthermore, Lightfoot and Able (2010) contended that previous microscopic studies (Keeling and Banttari, 1975; Van Caeseele and Grumbles, 1979; Keon and Hargreaves, 1983) used *P. teres* f. *maculata* isolates, arguing that these investigations predated the formal differentiation between *P. teres* f. *teres* (net form net blotch) and *P. teres* f. *maculata* (spot form net blotch). A re-examination of these papers, however, shows they consistently describe characteristic net-like lesions on the leaves, the defining symptoms of *P. teres* f. *teres*. This consistent description is why we chose these studies as foundational references for our microscopic investigation of *P. teres* f. *teres*.

Given the information above, we propose an infection model for the virulent *P. teres* f. *teres*–barley interaction. Within the first 12 hours post-inoculation (hpi), a germination tube emerges from the conidium and forms an appressoria-like structure that penetrates the epidermal cell. Inside the epidermal cell, an intracellular vesicle with appendages forms. The new intracellular vesicle has a dense, organelle-rich cytoplasm and its development is accompanied by a loss of epidermal cell membrane integrity (Keon and Hargreaves, 1983). The intracellular appendages penetrate the mesophyll layer, and intercellular mesophyll colonization is initiated. As infection progresses, the intracellular vesicle matures into a large, vacuolated structure (Keon and Hargreaves, 1983). By 48 hpi, intercellular mesophyll colonization continues, with hyphae growing in a form-fitting manner around mesophyll cells. PCD (nuclear breakdown) occurred behind the advancing hyphae (Danon et al. 2000; Domínguez and Cejudo, 2015). By 48 hpi intercellular hyphae in the mesophyll layer display hyphal stratification and by 72 hpi pericellular encasement. By 96 and subsequently 120 hpi, fungal hyphae have overtaken the mesophyll layer, colonizing much of the apoplastic space and leaving many mesophyll cells with ongoing PCD or completely collapsed.

Despite the early formation of specialized structures, our model reaffirms the classification of *P. teres* f*. teres* as a necrotroph, as its reproductive success remains tied to nutrient absorption from dying or dead host tissue resulting from PCD. These findings highlight the sophistication and nuances of fungal lifestyles. *P. teres* f. *teres’* delayed onset of PCD while building hyphal volume and strategically surrounding live cells appears to be a calculated necrotrophic manipulation of plant immunity. Furthermore, our observations of PCD trailing the hyphal front challenge generalizations that necrotrophs lack strategic complexity and add to growing evidence that they manipulate host immunity with precision (Kariyawasam et al. 2022).

### Comparative *Pyrenophora teres* f. *teres* infection timeline and volume analysis

After identifying and confirming *P. teres* f*. teres* infection structures, we analyzed the impact of VR1 and VR2 on the barley cultivar Rika at nine time points ranging from 4 to 120 hpi. Four specific isolates were used to evaluate and compare their effect on *in planta* colonization: #5*^vr1/vr2^* which lacks both *vr1* and *vr2*; #60*^VR1/vr2^*, which harbors only *VR1*; #72*^vr1/VR2^*, which harbors only *VR2*; and 6A*^VR1/VR2^*, which harbors both *VR1* and *VR2*. We observed no visible morphological or significant volumetric differences among isolates between 4 and 24 hpi; each successfully progressed through spore germination, appressoria-like structure formation, development of intracellular vesicles and appendages, and initiation of vertical intercellular hyphal growth in the mesophyll. Our findings indicate that the infection structures formed by 24 hpi are integral to *P. teres* f. *teres* infection and colonization. Additionally, their formation is not exclusive to *P. teres* f. *teres* isolates containing the *VR1* and/or *VR2* effector genes.

#### 48 to 72 hours post inoculation (hpi)

By both 48 hpi and 72 hpi, virulent isolates (#60*^VR1/vr2^*, #72*^vr1/VR2^* and 6A*^VR1/VR2^*) exhibited rapid intercellular colonization within the mesophyll that resulted in cytological features associated with PCD near older colonized regions. Conversely, #5*^vr1/vr2^*, exhibited limited mesophyll colonization. There was no significant difference in hyphal volume at 48 hpi and one significant difference at 72 hpi. We hypothesize that secretion of the *P. teres* f. *teres* effectors VR1 and/or VR2 promotes fungal growth in the mesophyll layer by targeting the susceptibility locus *Spt1.R* and facilitating the secretion of additional effectors into the mesophyll apoplast. *P. teres* f. *teres* establishes itself within the mesophyll before the induction of PCD, enabling modulation of the negative impact of the plant defense response.

#### 96 to 120 hours post inoculation (hpi)

The contribution of VR1 and VR2 became the most pronounced by 96 and 120 hpi. By 96 hpi, #5*^vr1/vr2^* continued to exhibit restricted mesophyll colonization and isolated instances of PCD but had hyphal volume that was not significantly different relative to isolates containing only one effector gene (#60*^VR1/vr2^* and #72*^vr1/VR2^*). However, 6A*^VR1/VR2^*, which contains both *VR1* and *VR2,* had visibly more *in planta* colonization, macroscopic lesions, and significantly more hyphal volume relative to all other isolates. These patterns continued at 120 hpi relative to #5*^vr1/vr2^* and #72*^vr1/VR2^*. Interestingly, #60*^VR1/vr2^* had visibly more mesophyll colonization, but did not have significantly different hyphal volume relative to #5*^vr1/vr2^*, #72*^vr1/VR2^* and 6A*^VR1/VR2^*. #60*^VR1/vr2^*’s hyphal volume was trending higher compared to both #72*^vr1/VR2^* and #5*^vr1/vr2^* and lower compared to 6A^VR1/VR2^.

Given the above information, we hypothesize a synergistic interaction between the two effector genes, *VR1* and *VR2*. Once initial mesophyll colonization was established, the presence of both effector genes enabled the pathogen to rapidly overwhelm host tissues, resulting in significantly larger hyphal volume at 96 and 120 hpi. Separately, *VR1* appeared to contribute more relative to *VR2* in colonization at later timepoints consistent with previously reported QTL effects at each locus (35% and 20% respectively; Shjerve et al. 2014).

Additionally, these findings suggest that 12, 48, and 96 hpi represent pivotal time points in the infection process, corresponding to the establishment of the pathogen in the mesophyll layer at 12 hpi, the induction of PCD in compatible interactions at 48 hpi, and the synergistic interactions of VR1 and VR2 resulting in large amounts of hyphal volume at 96 hpi. We propose that *P. teres* f. *teres* effector genes modulate plant defenses to facilitate accelerated accumulation of *in planta* volume in preparation for efficient nutrient absorption upon plant cell death. Increased fungal colonization results in a more robust induction of host PCD, which is macroscopically visible as expanded necrotic lesions. Ultimately, widespread host cell death provides the pathogen with the nutritional resources needed to sustain the *P. teres* f. *teres* polycyclic disease cycle.

## Conclusion

This study connects microscopic cellular events associated with pathogen infection to macroscopic leaf symptoms, while also characterizing the specific infection structures and compatible infection timeline for *P. teres* f. *teres* in barley. Additionally, this study shows that *VR1* and *VR2* together promote more rapid colonization of susceptible barley and identifies critical infection timepoints for disease development. Identifying these critical timepoints can inform the timing of fungicide application, potentially narrowing the window to maximize chemical efficacy and minimize environmental impact. Additionally, by clarifying the impact VR1 and VR2 have in driving colonization, this work lays the foundation for future effector characterization that can be used in crop improvement programs.

## Acknowledgements

This work was supported [in part] by the U.S. Department of Agriculture, Agricultural Research Service through CRIS project 3060-22000-051-000D and by the NSF/NIFA-AFRI Plant Biotic Interaction program of the National Science Foundation (award #1759030) and the USDA, National Institute of Food and Agriculture (award #2018-67014-28491).

The authors are grateful to Danielle Holmes for technical assistance, and Dr. Amber English at Imaris for technical software support.

## Disclaimer

Mention of trade names or commercial products in this publication is solely for the purpose of providing specific information and does not imply recommendation or endorsement by the U.S. Department of Agriculture. USDA is an equal opportunity provider and employer.

## Data availability

Sequencing information is available on NCBI (Accession numbers: SRX36034396, SRX36034398, and SRX36034397). Confocal images are available upon request.

## Author contributions

**Ashley C. Nelson:** Conceptualization, data curation, formal analysis, investigation, methodology, project administration, software, validation, visualization, writing – original draft, writing – review & editing. **Ryan M. Skiba:** data curation, formal analysis, investigation, methodology, validation, writing – review & editing. **Michele C. Malvestiti:** investigation, writing – review & editing. **Gayan Kariyawasam:** investigation, writing – review & editing. **Pawel Borowicz:** supervision, writing – review & editing. **Richard D. Horsley:** formal analysis, writing – review and editing. **Zhaohui Liu:** writing – review & editing. **Robert S. Brueggeman:** funding acquisition, writing – review & editing. **Timothy L. Friesen:** Conceptualization, funding acquisition, methodology, project administration, resources, supervision, writing – review & editing.

