## Supplemental Tables 1-3 for "*Pyrenophora teres* f. *teres* uses specialized infection structures to facilitate effector-driven promotion of *in planta* colonization on barley"

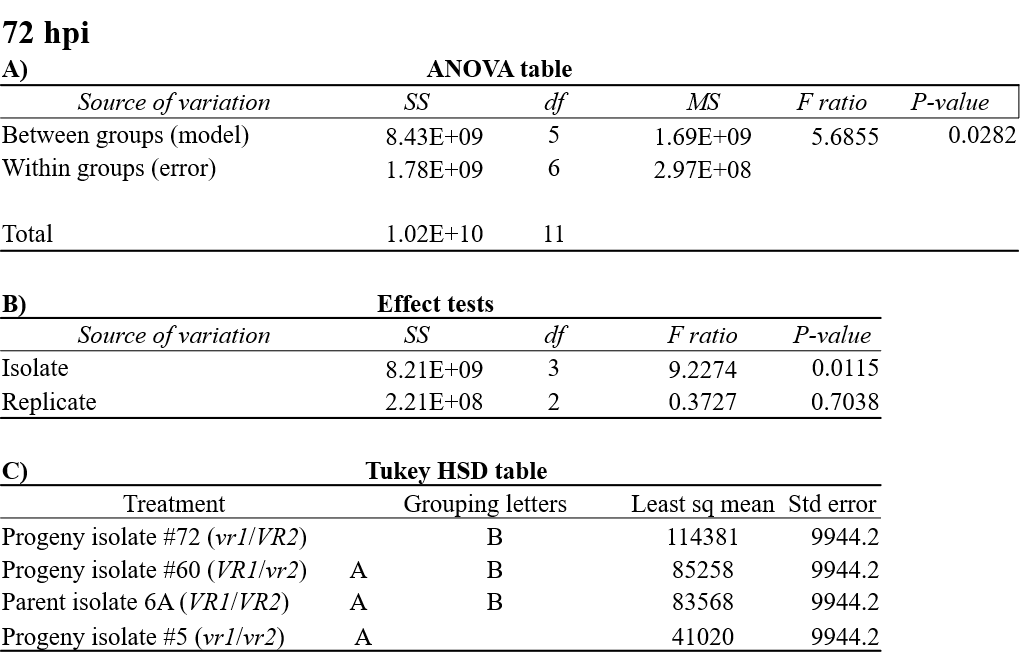

**Table S1: Statistical analysis of *Pyrenophora teres* f. *teres* isolates at 72 hours post inoculation (hpi)** **A.** ANOVA table for fungal volume (µm^3^) by isolate and replicate at 72 hpi. **B.**  Effect tests on isolate and replicate **C.** Tukey’s honestly significant difference (HSD) test for fungal volume (µm^3^) by isolate, and replicate following a significant ANOVA.

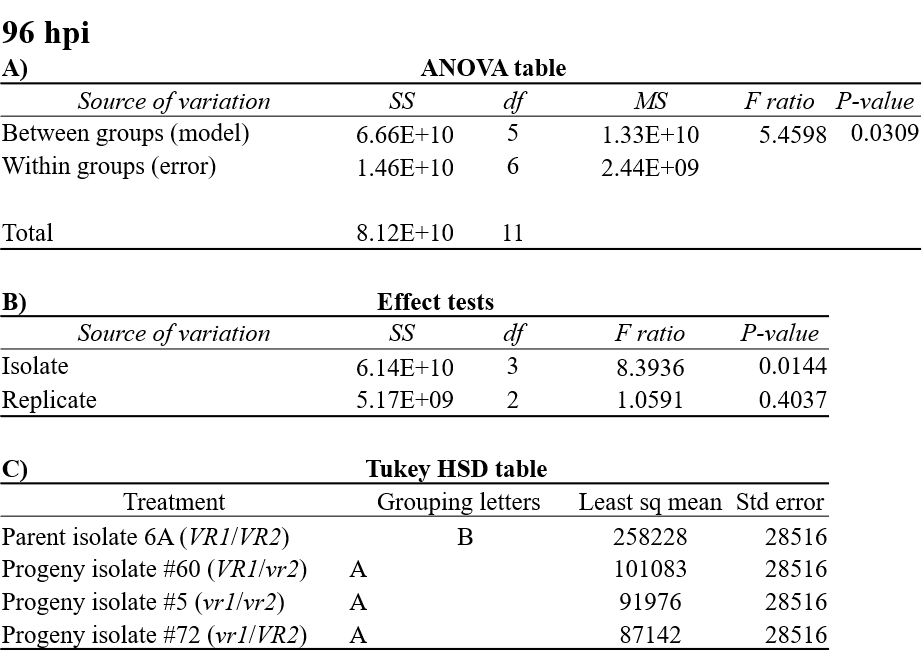

**Table S2: Statistical analysis of *Pyrenophora teres* f. *teres* isolates at 96 hours post inoculation (hpi)** **A.** ANOVA table for fungal volume (µm^3^) by isolate and replicate at 96 hpi. **B.**  Effect tests on isolate and replicate **C.** Tukey’s honestly significant difference (HSD) test for fungal volume (µm^3^) by isolate, and replicate following a significant ANOVA.

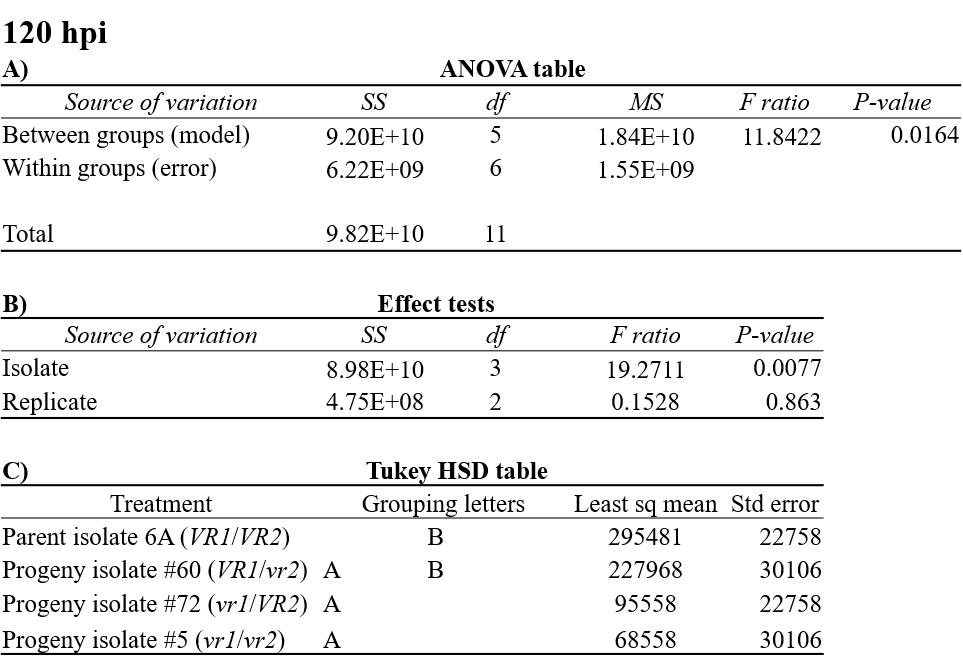

**Table S3: Statistical analysis of *Pyrenophora teres* f. *teres* isolates at 120 hours post inoculation (hpi)** **A.** ANOVA table for fungal volume (µm^3^) by isolate and replicate at 120 hpi. **B.**  Effect tests on isolate and replicate **C.** Tukey’s honestly significant difference (HSD) test for fungal volume (µm^3^) by isolate, and replicate following a significant ANOVA.
